# Fungi ligE-type *glutathione S-transferases* horizontally transferred into plants and a protozoan

**DOI:** 10.1101/2020.06.24.168203

**Authors:** Dan Li, Meng Zhang

## Abstract

Horizontal gene transfers (HGT) were considered as common evolution approaches for organisms. However, most HGT especially those HGT among distant species, like microbes to plants, were over-estimated because they were just identified based on the criteria of Blast searches and gene tree–species tree reconciliation. Recently, a ligE-type glutathione S-transferase (GST), *Fhb7-GST* was considered as an HGT from *Epichloë* to *Thinopyrum elongatum*. To confirm and clarify the occurring patterns of this HGT, homologue searches were conducted. Although *TeFhb7-GST* was not found in other plants, ligE-GSTs were found in not only plants but also an ameba protozoan. Additionally, *ligE-GSTs* were likely to horizontally transfer from fungi to other organisms. LigE-GSTs evolve in various fungi, but they only exist in some liverworts and green algae. Interestingly, all these *ligE-GST* genes in these plants share more than 90% similarities with that from fungus *Coniosporium apollinis*. More than that, the protozoan homologue from *Acanthamoeba castellanii* have 94.9% similiarity with that from *C. apollinis*. Actually, only a few substitutions were found in two homologues except a 111-bp lost in *A. castellanii ligE-GST*. All these results suggested HGT is an important evolutionary method for all organisms. Notably, natural HGT remind us to reevaluate the transgenic crops.

## Introduction

*Glutathione S-transferase* (GST, EC 2.5.1.18) genes are a supergene family and widely presented in probably all life forms [1]. GST enzymes majorly involved in the detoxication of a wide variety of endogenous and exogenous electrophiles by glutathione conjugation [2]. Also GSTs have several non-detoxification functions, for instance in plants, the non-enzyme carrier for intracellular transport and catalyzing the combination of anthocyanin and glutathione, and transported to vacuole by glutathione pump [2, 3]. Due to their multifunction characteristic, importance of existing in every living species examined and polymorphic, GSTs had been considered as valuable clinical candidates associated with disease phenotype in human and molecular breeding targets in crops [4, 5]. GSTs could divide into different clades among microbes, plants, animals and human depend on their similarities and functions [2-4, 6]. Furthermore, selective pressure is very strong in subfamilies of GSTs, probably resulting in functional diversifications of GSTs [7]. More than that, more and more new types of GSTs are continuing to be reported in various organisms, especially in plants [3, 8].

GSTs were considered have arisen from an ancient progenitor gene through both convergent and divergent pathways [9]. Probably, exon shuffling, gene duplication, alternative splicing, swapping, mutagenesis and other unknown mechanisms have led to considerable sequence diversification, functional heterogenicity and finally evolution of GSTs [7]. However, several reports found that horizontal gene transfer (HGT) was an important way to get useful GSTs for environmental adaption of all species, such as arthropod aerbivory [10].

Horizontal gene transfer (HGT) moves genetic material between or among lineages, and particularly has long been seen as a crucial process in the evolution of prokaryotic species [11, 12]. Currently, series methods, based on a combination of Blast searches and gene tree–species tree reconciliation, were developed to screen HGT genes from distantly related species (simply called as distant HGT, dHGT) [11, 13]. Commonly, most dHGT genes did not share high similarities of nucleotide sequences in distant species, and those dHGT from distant lineages, either evolved new classifications with new functions [14], or appear to be non-functional [15]. Hence, several researchers thought most dHGT genes were over-estimated, especially those genes transferred among eukaryotes [11, 16]. A more convincible method is needed to evaluate dHGT.

Recently, a *Thinopyrum elongatum GST* gene, which confers broad resistance to *Fusarium* species, was gained via HGT from endophytic *Epichloë* species [17]. *TeFhb7-GST* from *Thinopyrum elongatum* has 97% similarity with that from *Epichloë* fungi. Moreover, *TeFhb7-GST* encodes a ligE-type GST, which belongs to fungal GTE (glutathione transferase esterase-related) subfamily wherein all members has a ligE domain [17, 18]. Except in *Thinopyrum* species, no homologues of *TeFhb7-GST* were found in entire plant kingdom according to NR database [17]. It’s a solid proof of dHGT that transferred from fungi to plants, and told us that dHGT is essential for plants to improve pathogen resistance.

According to these results, HGT is definitely one reason for the expansion and multiclassification of GSTs. But it’s still confused that whether dHGT of *TeFhb7-GST* and ligE-type GST genes are either common or coincidence in organisms. Therefore, a deep study of homologue searching of ligE-type *TeFhb7-GST* were conducted. Finally, a protozoa ligE-type GST was identified in *Acanthamoeba castellanii* and certificated that *AcGTE* was a dHGT transferred from fungi to both plants and *A. castellanii*. All these results showed the solid evidence of dHGT, but the meaning need to be clarified in protozoa and these organisms.

## Results

### Homologues of *TeFhb7-GST* are all ligE-GST encoding genes

To find the homologues of *TeFhb7-GST*, genome databases of Phytozome V12 [19], OneKP (https://db.cngb.org/blast/) and ConGenIE [20] were employed. Finally, series of homologues were found to have considerable similarities (about 60%) with *TeFhb7-GST* in plants, fungi and even in a protozoan, (Figure 1). Although these genes shared only ∼60% similarities with *TeFhb7-GST*, but they all encoded ligE-type GSTs that similar with *Phanerochaete chrysosporium* GTE1 which is the firstly identified LigE-GST, suggesting that HGT of ligE-GST is not a coincidence from *Epichloë* fungi to *Thinopyrum* plant (Figure 2).

**Figure 1.**
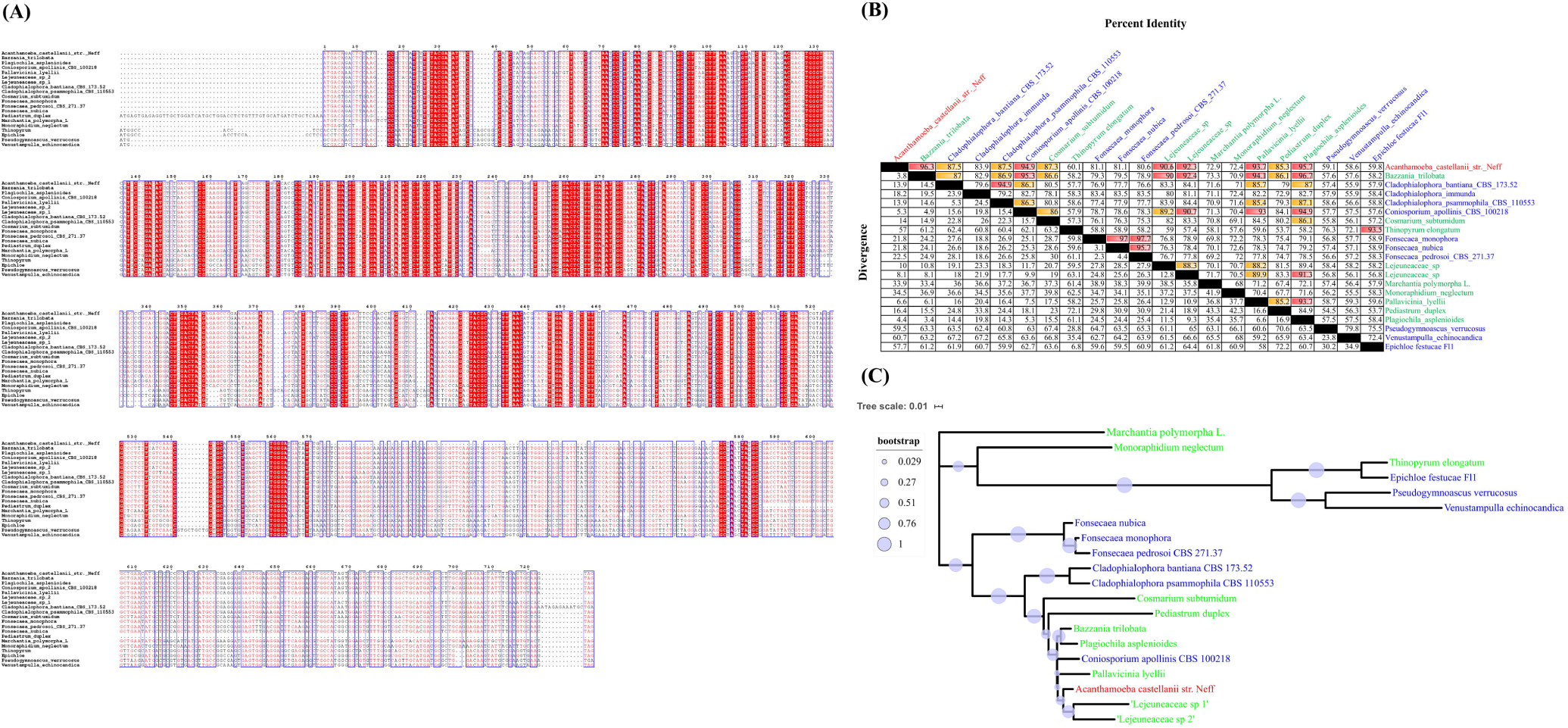
*Fhb7-GST* homologues. The coding sequences of *Thinopyrum elongatum* Fhb7-GST homologues were retrieved from variety species. They were aligned with MAFFT (A), the similarities and divergence of each two were displayed (B). And the phylogenic of nucleotides were made by MEGA X using the Maximum Likelihood method and General Time Reversible mode (C).

**Figure 2.**
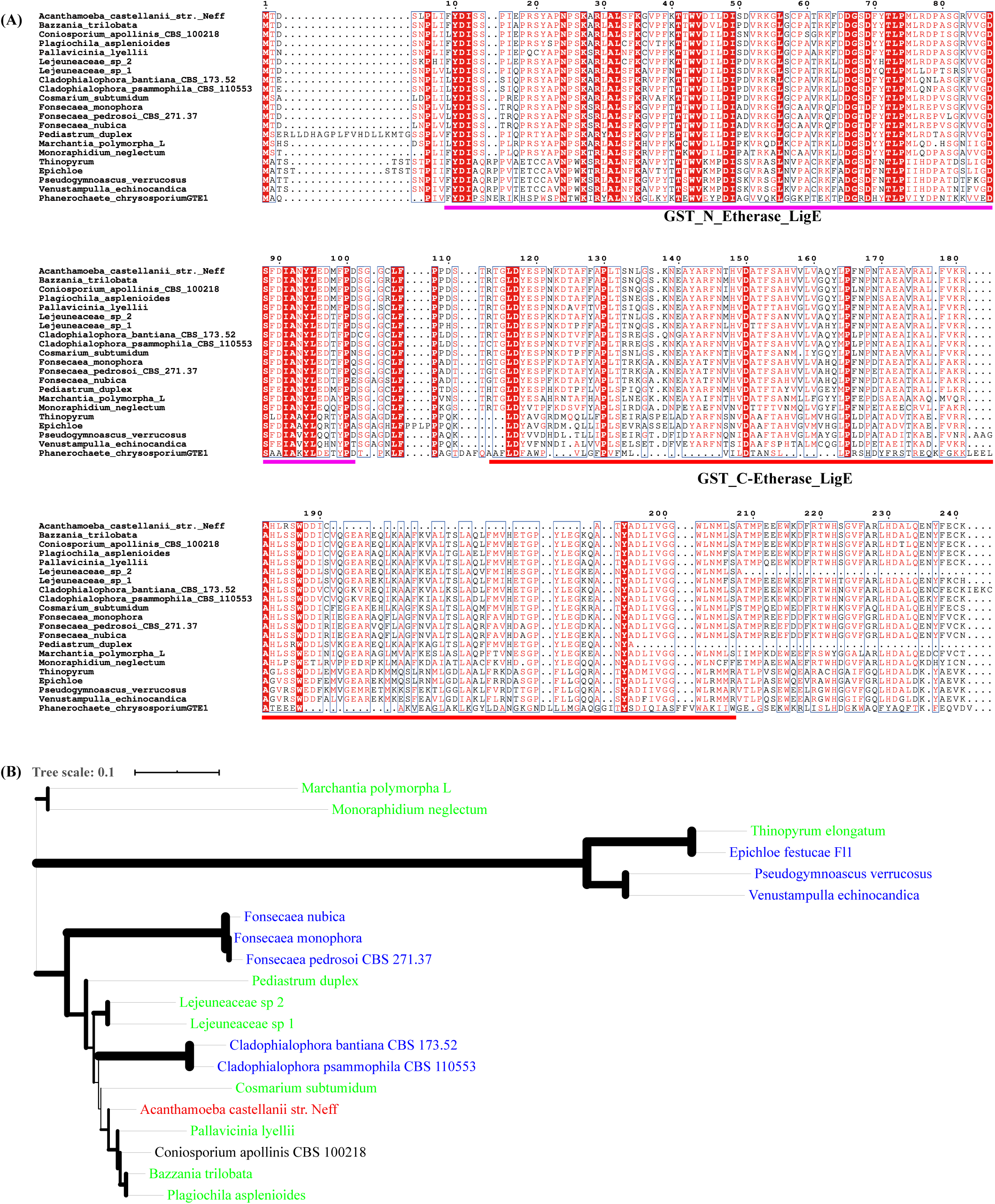
Fhb7-GSTs all encode LigE-GSTs. The protein sequences of these GSTs were retrieved from variety species. They were aligned with MAFFT (A). And the phylogenic of nucleotides were made by MEGA X using the Neighbor-Joining method and Jones-Taylor-Thornton mode (B).

### Homologues of *TeFhb7-GST* exists in fungi, Marchantiophyta, green algae and amoebae

Although these ligE-GSTs presented in several plants, except *Thinopyrum*, all these plants were either Marchantiophyta (*Bazzania trilobata, Lejeuneaceae, Marchantia polymorpha* L., *Pallavicinia lyellii* and *Plagiochila asplenioides*) or green algae (*Cosmarium subtumidum, Monoraphidium neglectum, Pediastrum duplex*) (Figure 1). The ligE-type GSTs were not found in other plants.

The gene in *Monoraphidium neglectum* was the closest relative of *Fhb7-GSTs*. But homologues from other green algae were closer to fungi and several Marchantiophyta, suggesting these genes were obtained from HGT.

As the previous report, *TeFhb7-GST* shared 91%—97% similarities with those from several *Epichloë* species, such as *E. aotearoae*, suggesting *Fhb7-GSTs* were polymorphically evolved in *Epichloë* fungi *[17]*. Besides *Pseudogymnoascus verrucosus* (∼75% similarity), a *TeFhb7-GST* homologue (also ∼75% similarity) was also found in *Venustampulla echinocandica*, which produces extracts with antifungal activity against *Aspergillus fumigatus* and *Candida albicans* [21, 22]. And *PvFhb7-GST* and *VeFhb7-GST* has 79.8% similarity (Figure 1b). But in other fungi, ligE-type GST encoding genes have no more than 60% similarities with *Fhb7-GSTs*, suggesting ligE-type GST have multiple functions. Indeed, except *TeFhb7-GST*, none ligE-type GST was functionally characterized to date [17].

Interestingly, a ligE-GST homologue gene was found in a protozoan, *Acanthamoeba castellanii*. The 2.9% proteins of *Ac. castellanii* were previously considered to be obtained by HGT *[23]*. Now, the *Ac. castellanii ligE-GST* confirmed that it was indeed a foreign DNA and obtained from fungi, maybe from *Coniosporium apollinis*.

### *Ac. castellanii ligE-GST* is an HGT gene probably transferred from *Coniosporium apollinis*

Amongst these ligE-GSTs, *Ac. castellanii ligE-GST* is another HGT gene and probably transferred from *Coniosporium apollinis* fungi into several plants. *Ac. castellanii ligE-GST* showed more than 93% similarities with those homologues in liverworts and algae, including *Bazzania trilobata* (96.3%), *Pallavicinia lyellii* (93.7%) and *Plagiochila asplenioides* (95.7%). Actually, all these genes were highly similar with that in *Coniosporium apollinis*. Indeed, without the lost 111 bp in the middle, *Ac. castellanii Fhb7-GST* has only few substitutes with other homologues in these organisms. It indicated that all these homologues were HGT genes and might origin from *Coniosporium apollinis*.

Additionally, there were several homologues sharing more than 85% similarities from two *Lejeuneaceae* liverworts (90.6% and 92.3%), *Cladophialophora* fungi (83.9%∼87.5%) and *Pediastrum duplex* (green algae, 85.3%). Since the similarities of these homologue genes were quite different from species tree, they are probably all obtained via HGT. That’s to say, several fungi were missing to contain similar genes with those in *Lejeuneaceae* liverworts and green algae *Pediastrum duplex*.

Due to the limited information of functional characterization of ligE GSTs, it’s still unknown what the meaning of *AcFhb7-GST* is for the pathogenic *Acanthamoeba castellanii*, and we are still wonder how and why these organisms obtained these HGT genes.

## Discussion

The ligE-type GSTs were firstly found in fungus *Phanerochaete chrysosporium*, then were certificated to be widely present in various fungi [17, 18]. The ligE GSTs possess the specific GST-N and GST-C domains of a bacterial GST, named as ligE protein in *Sphingomonas paucimobilis*. Perhaps, fungal ligE-type GSTs were evolved from bacterial ligE proteins.

Although these GSTs from fungi and bacterial are related, their functions are quite different [18]. Bacterial LigE catalyzes the reductive cleavage of the β-aryl ether linkage of guaiacylglycerol β-O-4-methylumbelliferone to produce β-hydroxypropiovanillone and 4-methylumbelliferone in vitro [24]. Whereas, the *P. chrysosporium* ligE-type GSTs seemed not to be active using this substrate [18].

Previously, ligE-type GSTs were never identified in plants. Interestingly, a *Fhb7*-*GST* gene was confirmed to encode a ligE-type GST in *Thinopyrum elongatum*, which was a higher plant and closely related to wheat. Firstly, Wang et al. found that TeFhb7-GST could detoxify trichothecenes through de-epoxidation to confer broad resistance to *Fusarium* species, which cause food toxins, currently devastates wheat production worldwide [17]. Furthermore, the nucleotide sequence of *TeFhb7-GST* shares 97% similarities with homologues from fungi *Epichloë* species. It seems that *TeFhb7-GST* was an HGT gene that transferred among distant species. This result solidly certificated that HGT is a real phenomenon in lives.

The HGT of *TeFhb7-GST* is beneficial for *Thinopyrum elongatum* to resist the infections from harmful *Fusarium* species actually. But it’s strange that only *Thinopyrum* species obtained the detoxification *GST* gene whereas *Fusarium* head blight is a heavy fungal disease for wheat, barley and other small grain cereals. And *Epichloë* species are endophytic fungi commonly living in most C3 grasses including poaceae plants, such as wheat [25, 26]. Since no any similar genes were found in other plants, the HGT of *TeFhb7-GST* seemed to be a coincidence.

However, according to our results, HGT of *ligE-type GSTs* massively occurred among distant species. Several nucleotide sequences from liverworts, algae and even a protozoan, share about 60% similarities with *TeFhb7-GST*. Phylogenic analysis also indicated these genes encode ligE-type GST proteins. However, these proteins were not likely evolved. In many liverworts, there were no such GSTs according to reported genomes. Moreover, some of them share 95% similarities with each other, for instance, homologues in *Coniosporium apollinis* (fungus), *Bazzania trilobata* (liverwort) and *Ac. Castellanii* (protozoan). The most interesting is that it’s extremely similar between *Ac. Castellanii ligE-GST* and the other two homologues. These results suggested HGT of *ligE-type GSTs* was a common phenomenon, between fungi and not only plants but also animals. It seems that the frequency of occurrence of HGT is not low as previous thought, thus gene drifts are likely important adaption strategy for various organisms.

Since foreign genes are not unique for organisms, we might reconsider and reevaluate the applications of genetically modified crops. But, it is necessary to develop novel gene transformation technologies by clarifying the transfer mechanism of HGT, making sure only the essential nucleotide fragments were inserted in proper sites.

All these results showed that genes were actually horizontally transferred among distant species, but we should be more cautious to identify HGT. More convincible HGT are more helpful for us to understand the evolution of organisms. Finally, I hope the studies of distant HGT could help ordinary people and government rethink the supervising and applying of transgenic technologies in crop breeding.

## Materials and Methods

### Homologues searching and sequences obtaining

*Thinopyrum elongatum Fhb7-GST* was get from the work of Wang et al. 2020 [17], and then the nucleotide sequence was used to blast against Nr database to get the sequences of homologues from *Epichloë, Pseudogymnoascus verrucosus* and *Venustampulla echinocandica*. Then, the three nucleotides were used to blast against Phytozome V12, and only the homologue of *M. polymorpha* was obtained. Subsequently, *Marchantia polymorpha GST* was homologous searched in OneKP database, and other homologues were obtained. Also, these homologues were blasted against gymnosperm genomic datasets, such as ConGenIE, but there was no any similar sequence.

### Sequence alignment and phylogenic tree construction

All the nucleotides and proteins were aligned by MAFFT server at EBI with default settings, and then colored by ESPript 3.0. Phylgenic trees were constructed by MEGA X ver. 10.05, and visualized by iTOL (https://itol.embl.de/upload.cgi). For nucleotides, Maximum Likelihood method and General Time Reversible mode were used; while Neighbor-Joining method and Jones-Taylor-Thornton mode were used to construct the phylogenic tree. For each tree, 1000 bootstrap replications were applied.

## Compliance and ethics

The author(s) declare that they have no conflict of interest.

